# The influence of the supramolecular complex PRE -1 on permeability of membranes of mitochondria

**DOI:** 10.1101/300855

**Authors:** Kamila Mardieva, Munira Burkhanova

## Abstract

The change in the state of the mitochondrial pore of nonspecific conductivity plays a key role in the processes of triggering and regulating apoptosis. Therefore, the study of mechanisms of opening and regulating the state of these pores can have a great importance to find the ways of correcting many mitochondrial dysfunctions, including those which are associated with age-related changes. In our work, we have done a comparative analysis for swelling of brain mitochondria of adult (9 months) and mature rats (12-15 months).

## 1 Introduction

Mitochondria have a key role in the processes of apoptosis and necrosis, so it is very likely that the disturbance of mitochondrial functions, observed with aging is associated with a violation of the regulation of programmed cell death. Since it is known that the aging of many types of cells is associated with a change in their sensitivity to apoptosis. Therefore, there are reasons to assume that the mitochondrial dysfunction observed with aging is associated with a violation of the regulation of programmed cell death.

The discovery of permeability transition pore (PTP) on the inner mitochondrial membrane leads to: the swelling of these organelles (1), the dissociation of oxidative phosphorylation(2), the decrease in the synthesis of ATP molecules(3), which can be extremely detrimental to the condition of individual cells, and of whole tissues and organs, and ultimately of the organism as a whole. According to the main hypothesis, this pore is a mega-channel consisting of an adenyl translocase that underwent a pronounced conformational change that interacts with other proteins of the inner membrane. The study of the mechanisms of opening and regulation of the discovery of these pores can be of great importance for finding ways to correct many mitochondrial dysfunctions, including those associated with age-related changes.

## 2 Analysis

Quite often one compares the indicators of young and very mature, old organisms, while the processes leading to aging run at much earlier stages of growing up the body. Therefore, an approach that compares the indicators of young and mature but still fully functioning organisms can be considered more promising. In addition, prevention and correction of disturbances at the early stages of the development of aging processes is more effective, rather than treating the already existing dramatic changes. This approach will prolong the period of full-fledged vital activity and, ultimately, will ensure the state of “healthy old age”.

The purpose of our research is to study the effect of the supramolecular complex PRE-1 on the permeability of mitochondrial membranes of brain of rats of different ages.

Male white mongrel rats are used in the study. They have 3 different age category: 9, 12, 15 months.

In order to assess the effect of the supramolecular complex PRE-1 on the permeability of mitochondrial membranes, experiments were performed in vivo, administered intra-abdominally at a concentration of 30 mg / kg 1 time per day for 5 days.

Mitochondria were isolated by differential centrifugation [1] in a medium containing 320 mM sucrose, 10 mM Tris-HCl and 0.5 mM EDTA or 0.5 mM EGTA, 0.2% BSA, pH 7.4. The isolated brain was crushed, purified from blood vessels and destroyed in a tenfold volume medium, using a glass homogenizer. The homogenate was centrifuged at 2000 g for 3 min, the pellet was removed and the supernatant was centrifuged again to remove the nuclei and the damaged cells more completely. Mitochondria were precipitated by centrifugation at 12,500 g for 10 min at 4°C. The precipitate (1.5 ml) was mixed with 3% percol (1 ml), then the mitochondria were purified in a gradient of percol (10-24%), precipitating the non-synaptic mitochondria at 31,500 g for 10 min. The mitochondrial precipitate was washed with a release medium without EDTA, EGTA and BSA (11,500 g, 10 min) and suspended in the same medium.

*The protein content* was determined colorimetrically according to the biuret method [2] using bovine serum albumin as the standard.

*Swelling of rat mitochondria* was determined from the change in light scattering, at a wavelength of 540 nm, on an Agilent Technologies-Cary 60 spectrophotometer in an open cell (3 ml volume) with vigorous stirring at 25°C [3]. In an incubation medium containing 10 mM Tris-HCl, 120 mM KCl, 5 mM glutamate, 1 mM malate, and 2.5 mM KH2PO4.

## 3 Results and discussions

*In vivo* experiments it was found that in the absence of Ca^2+^ in the mitochondrial environment old rats are not swollen, and the sensitivity of mitochondrial membranes to the Ca^2+^ load depends on the age of the animal: for 9- and 12-month-old rats, controlled swelling is caused by the addition of Ca at a concentration of 40 *μM*, whereas for mature rats (15 months) is caused by the addition of already 20 *μM* Ca. This means that with the increase of the age, mitochondria become more sensitive to the effects of calcium ions.

Calcium stimulated swelling by 95 - 98% was blocked by a specific inhibitor of PTP cyclosporinA, which indicates the main role of this pore in the process of swelling of mitochondria. It is likely that the change with age of PTP sensitivity to calcium loading largely determines the changes in tissue sensitivity underlying cell damage during aging. The obtained results coincide with the results of other researchers who testify that the sensitivity of mitochondria to activation of ptp increases in aging, which is manifested in a decrease in the threshold concentration of Ca^2+^, which initiates the discovery of PTP [3].

In our work, a comparative analysis of the swelling of adult brain mitochondria (9 months) and mature rats (12-15 months) was performed. The load of mitochondria with calcium, the amount of which corresponded to the threshold concentrations, caused the acceleration of mitochondrial swelling in mature rats 4-5 times compared to the mitochondria of adult rats (see Figure 1). When comparing the half-periods of maximum swelling of mitochondria in adults and mature rats, reflecting the kinetics and confirming this conclusion, it was found that in mature rats this figure was reduced by 4 - 5.7 times. An increase in the rate of Ca^2+^ -induced swelling in older rats (12 and 15 months) resulted in a decrease in the time for complete swelling of the mitochondria-in 2 and 4 times in 12- and 15-month-old rats, correspondingly, in comparison with the indices of 9-month-old animals.

**Figure 1:**
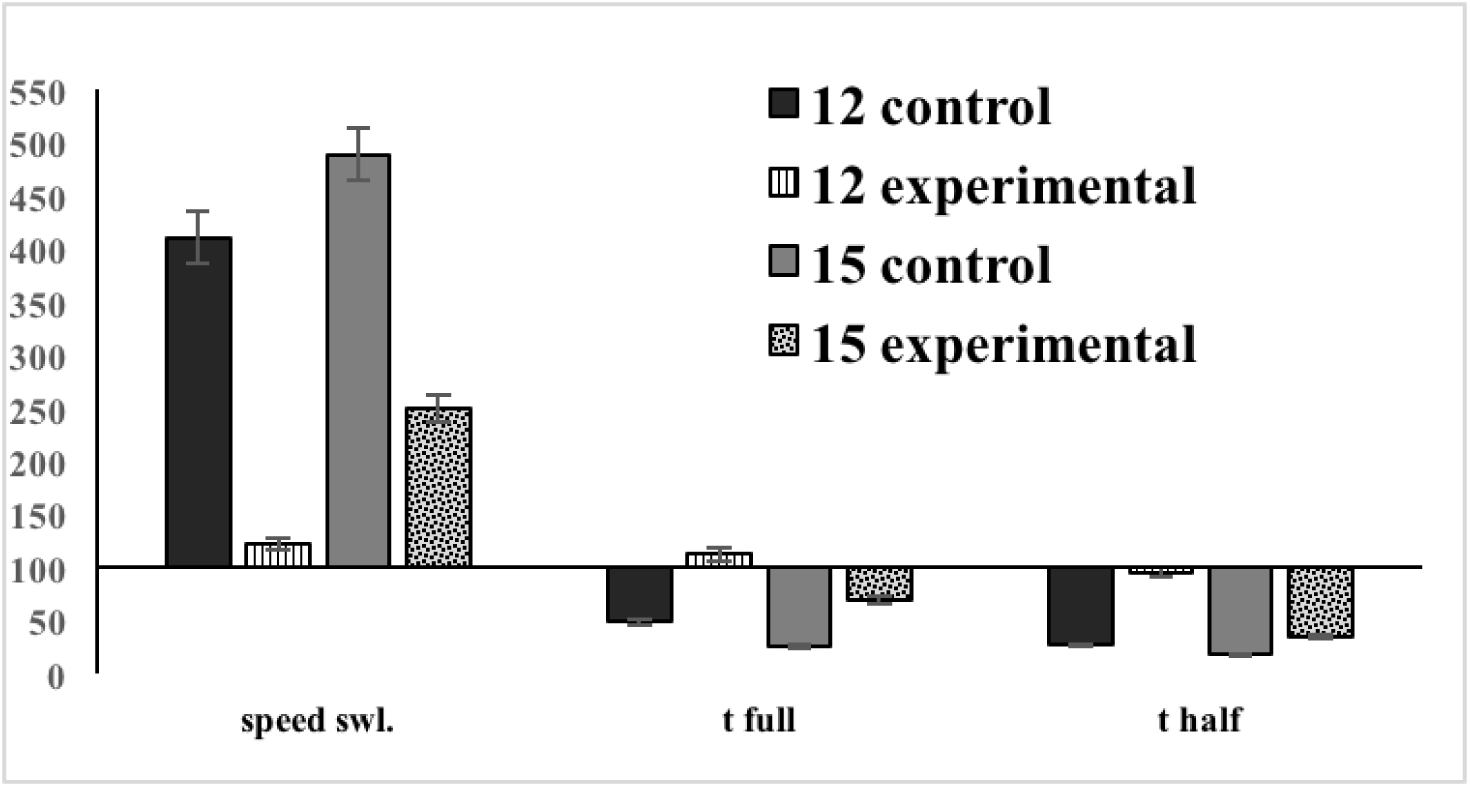
Age-related changes in the permeability of mitochondrial membranes (100% for the control group of 9-month-old animals).

It should be noted that in 9-month rats the kinetic pattern of swelling was significantly different from other ages: swelling occurred in 3 stages, with each subsequent stage proceeding at a faster rate than the previous one.

*In vivo* experiments, we found that the PRE I supramolecular complex (SMC), injected intraperitoneally at a concentration of 30 mg / kg 1 time per day during 5 days, had a significant effect on Ca-induced permeability of mitochondrial membranes, the orientation of which depends on the age of the treated rats (see Table 1).

**Table 1:** Effect of SMC PRE - I on Ca^2+^ -induced swelling of energized mitochondria

| | Age<br>(month) | $\text{Ca}^{2+}$ injection<br>($\mu\text{M}$) | Swelling<br>(opt. dens. per min.) | t full<br>(min) | t half<br>(min) |
| --- | --- | --- | --- | --- | --- |
| 9 | control | 40 | $0.18 \pm 0.02$ | $6.17 \pm 0.06$ | $4.3 \pm 0.07$ |
| | experimental | | $0.36 \pm 0.07$ | $6.17 \pm 0.06$ | $2.7 \pm 0.07$ |
| 12 | control | 40 | $0.74 \pm 0.01$ | $3.02 \pm 0.03$ | $1.14 \pm 0.02$ |
| | experimental | | $0.22 \pm 0.01$ | $6.92 \pm 0.08$ | $4.1 \pm 0.04$ |
| 15 | control | 20 | $0.88 \pm 0.03$ | $1.56 \pm 0.01$ | $0.76 \pm 0.01$ |
| | experimental | | $0.45 \pm 0.01$ | $4.31 \pm 0.02$ | $1.47 \pm 0.01$ |

Since both in the control group and in the group of treated rats, mitochondrial swelling was almost completely inhibited by a specific inhibitor of the mitochondrial pore of nonspecific conductivity with cyclosporin A, it can be safely assumed that the change in membrane permeability is directly related to the change in the state of the given pore.

In the youngest group (9 months), the 5-day administration of the drug has a negative effect, increasing the sensitivity of mitochondria to ions of Ca^2+^: in treated animals the speed of Ca^2+^ assimilability increases 2 times and and the half-maximal swelling time is reduced 2 times. However, for older rats (12-15 months), for which the control was characterized by a significant increase (by 4 or more times) in the swelling rate and a critical reduction in both the full and half-maximal swelling time, PRE - I had a pronounced corrective effect and characteristics of the mitochondria of 12-15 month old animals that received PreI, approached the control values of 9 month old animals. Therefore, PRE-I is a promising platform for creating medicines aimed at correcting age-related changes.

